# Kernel-Based Style Transfer Mapping for Cross-Subject Biological Signal Classification

**DOI:** 10.1101/2025.09.01.673597

**Authors:** Takayuki Hoshino, Suguru Kanoga, Atsushi Aoyama

## Abstract

Owing to pronounced inter-individual variability in biological signals, transfer learning has emerged as a widely used strategy to reduce calibration requirements for new users. Among the various approaches, style transfer mapping (STM) is distinguished by its ability to align the data distribution of target users directly with that of source users, thereby enabling the reuse of pretrained classifiers without retraining. However, STM is constrained by its inherent assumption of linear relationships between the target and source domains, which is often unrealistic given the complex and nonlinear nature of biological signals. To address this problem, we propose a novel kernel-based STM (k-STM) method that incorporates nonlinear kernel functions to capture complex inter-user variability more effectively. We evaluated the proposed k-STM method using four publicly available electromyogram and electroencephalogram datasets: MyoDataset, NinaProDB5, OpenBMI, and BCI Competition IV-2a, and four backbone classifiers: logistic regression, support vector machine, random forest, and multilayer perceptron. The experimental results demonstrate that k-STM significantly improves classification accuracy over the original STM, particularly when nonlinear kernels are employed. Statistical analyses confirm the superiority of the proposed approach across diverse datasets and classifiers. Furthermore, we examined the impact of the number of prototypes (i.e., mapping destinations) on the performance of both STM and k-STM, and found that using fewer prototypes than previously recommended can yield higher classification accuracy. Overall, the proposed k-STM constitutes a versatile, efficient, and robust nonlinear transfer-learning framework, enhancing its practical applicability in cross-subject biosignal classification.

## 1 Introduction

Decoding user intentions from biological signals using machine learning is crucial for the development of next-generation human–computer interfaces (HCIs) [1, 2]. For instance, the decoding of surface electromyogram (sEMG) and electroencephalogram (EEG) signals has been applied in various applications, including the control of prosthetic limbs [3, 4] and wheelchairs [5, 6]. Achieving high classification accuracy is essential for practical and reliable HCIs because misclassification can lead to incorrect or even harmful device actions. Traditionally, classifiers have been trained on a per-user basis to ensure high accuracy owing to the substantial inter-subject variability in biological signals [3–6]. However, collecting sufficient data for each user is time-consuming and burdensome, posing a major challenge for real-world deployment.

Transfer learning has emerged as a powerful approach to address this issue [7]. By mitigating variabilities in data distribution, transfer learning enables the effective utilization of previously collected data from other users and pretrained classifiers [8, 9]. Consequently, transfer learning significantly reduces the data-collection burden and improves classification accuracy, helping overcome key obstacles to practical HCI implementation.

Numerous transfer learning approaches have been proposed for sEMG and EEG signal classification [8,9]. Among these, style transfer mapping (STM) has emerged as a practical technique for adapting pretrained classifiers to new (target) users. Originally developed for handwritten character recognition [10], STM was later applied to EEG signals [11] and subsequently extended to sEMG signals [12]. Figure 1 provides an overview of STM. A classifier is first trained using data from a previously recorded (source) user (denoted by ‘ º’ markers). When this classifier is applied directly to data from a new (target) user (represented by ‘ ×’ markers), inter-subject variability significantly degrades classification accuracy. STM addresses this issue by learning a mapping function that projects the data of the target user onto the data distribution of the source user. This mapping mitigates the distribution shifts caused by individual variabilities, enabling the direct reuse of the pretrained source classifier for the target user.

**Figure 1:**
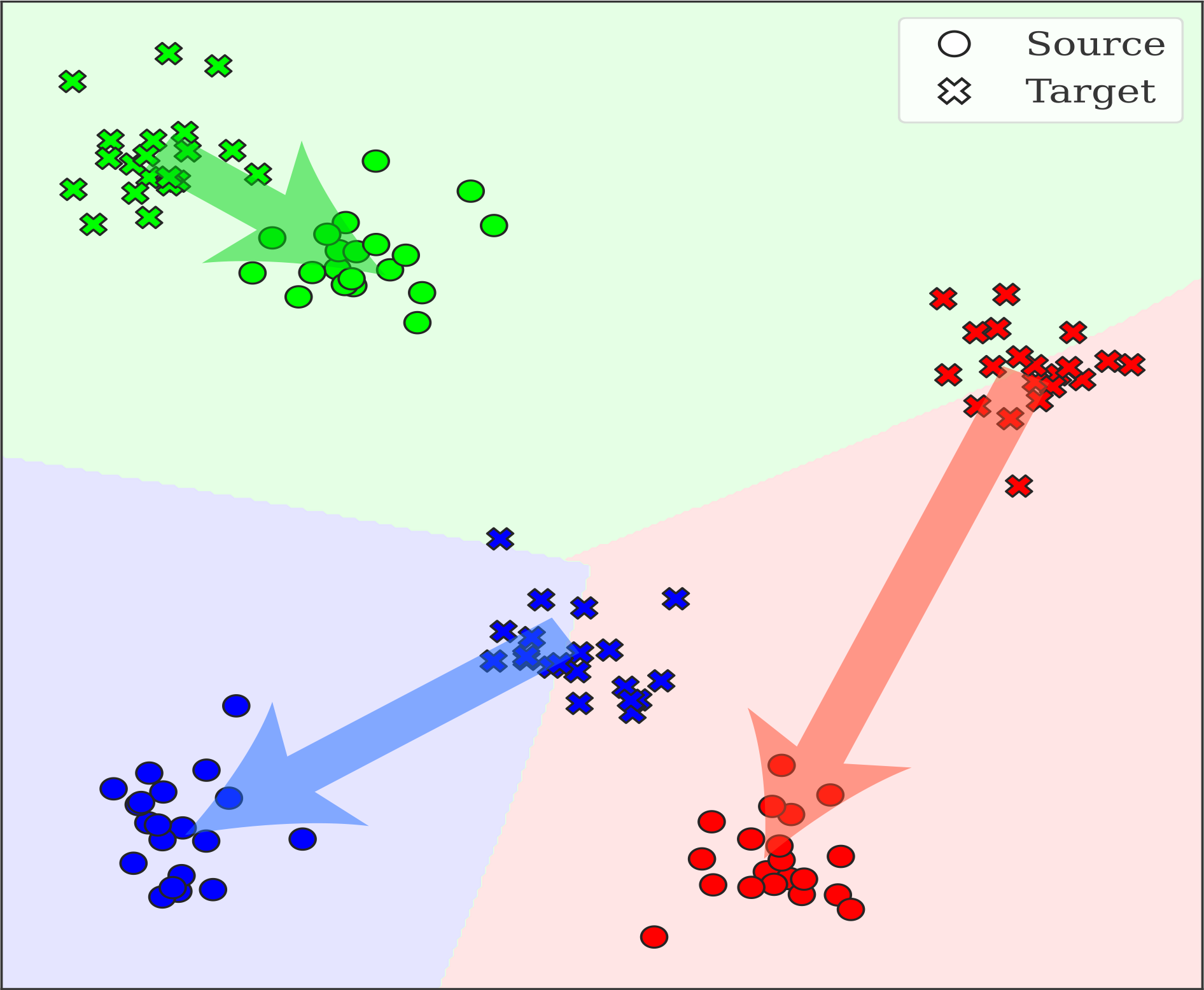
Overview of style transfer mapping (STM). Source user data (º) and target user data (×) are shown. Background colors indicate decision regions learned from the source user’s data. Because of inter-subject variability, the target user’s data are shifted and misaligned, preventing accurate classification by the source-trained classifier. STM addresses this issue by mapping the target data along the directions indicated by arrows, thereby aligning them with the source data distribution and enabling accurate classification using the source classifier.

Compared with other transfer learning methods, STM offers two key advantages: (1) direct reuse of a pretrained source classifier without retraining, and (2) compatibility with any classifier architecture. Regarding the first advantage, although other mapping-based methods such as transfer component analysis [13] and correlation alignment [14] have been proposed, these approaches require mapping both the source and target data into a common subspace. Consequently, they require the training of a new classifier from scratch on the transformed data, resulting in high computational costs for each new user. By contrast, STM maps only the target data, allowing the source classifier to be reused without retraining. The second advantage is the flexibility of the model. Parameter-based transfer learning methods—such as covariate shift adaptation [15], fine-tuning [16], domain-adversarial neural networks [17], and Deep CORAL [18]—permit reuse of source classifiers but are inherently tied to specific model architectures. Because the optimal machine learning model often varies across individuals and tasks, this dependency limits its practical generalizability. By contrast, STM is classifier-agnostic and universally applicable to existing pretrained classifiers, providing flexibility in selecting the most suitable model for each application. Together, these advantages highlight the practical versatility of STM, making it a strong candidate for transfer learning in biological signal classification tasks.

Recently, several STM-based improvements have been proposed [19–23]. Kanoga et al. introduced a method that applied STM prior to the fine-tuning of convolutional neural networks, demonstrating its effectiveness as a preprocessing step for subsequent transfer learning methods [19]. Niu et al. proposed combining STM with Euclidean alignment, showing that STM can also serve as an effective postprocessing step [20]. Ran et al. developed an instance-selection approach that applies the STM only to selected samples [21]. Chen et al. introduced a novel regularization technique that enables STM to perform well even with limited data [22]. Huang et al. proposed a strategy for fusing data from multiple source users into a smaller set of representative domains using STM [23]. These advancements highlight the versatility and strong potential of STM to enhance classifier adaptability across diverse domains and datasets.

Although several advanced STM-based methods have been proposed [11, 12, 19–23], STM has an inherent limitation in that it relies on a linear mapping function, and therefore cannot adequately capture the complex nonlinear characteristics associated with individual variability in biological signals. This is a significant drawback because biological signals, such as sEMG and EEG, inherently exhibit nonlinear dynamics arising from multiple physiological and cognitive factors. For example, sEMG signals are influenced by age- and sex-related differences in muscle strength [24] as well as by muscle fatigue [25]. Similarly, EEG signals are affected by genetic factors related to the brain structure [26] and fluctuating attention levels during task execution [27]. Given these properties, restricting STM to linear mappings is overly restrictive and often leads to underfitting. As illustrated in Figure 2(a) and (b), STM performs well under linear relationships but fails to capture nonlinear relationships. Therefore, extending the current linear STM to nonlinear forms is more suitable for the characteristics of biological signals and is expected to enable more accurate transfer learning.

**Figure 2:**
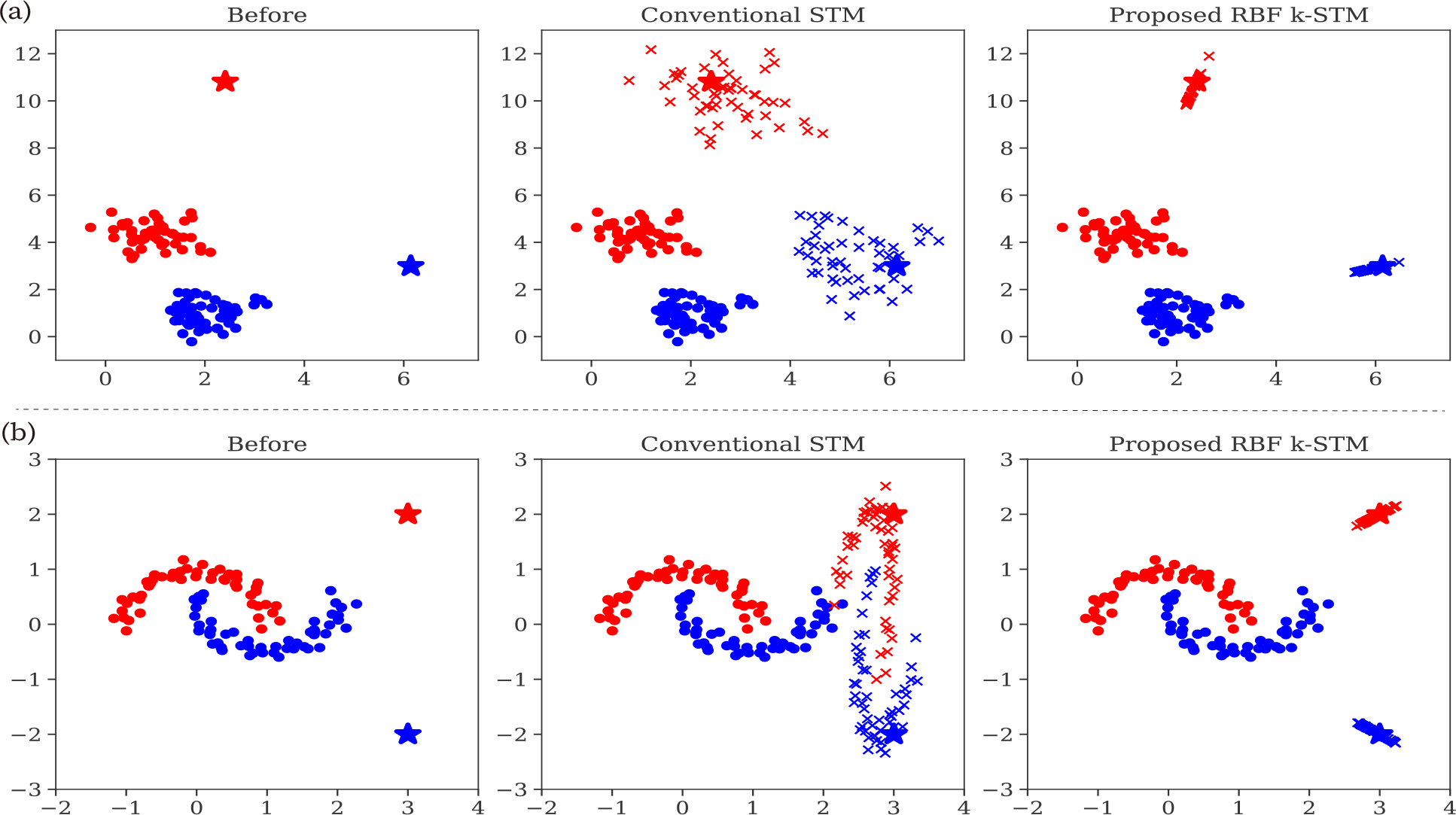
Examples illustrating STM performance under (a) linear and (b) nonlinear relationships. The left column shows the original distributions of source and target data. The middle column depicts the application of conventional STM, while the right column shows RBF-based k-STM. Circle markers (º) represent original target samples, which are mapped towards source-domain prototypes indicated by stars (⋆). Cross markers (×) denote the positions of the target data after mapping. Conventional STM fails to capture nonlinear mappings effectively, whereas RBF-based k-STM successfully handles both linear and nonlinear relationships.

To address this problem, we propose a novel kernel-based STM (k-STM), which extends the conventional STM by employing kernel methods to enable nonlinear transformations. By introducing nonlinear mappings, k-STM achieves an accurate transfer even when the relationship between the source and target domains is nonlinear, as illustrated in Figure 2(b). To evaluate the effectiveness of the k-STM, we conducted experiments on four publicly available sEMG and EEG datasets: MyoDataset [28], NinaProDB5 [29], OpenBMI [30], and BCI Competition IV-2a (BCICIV2a) [31]. We implemented the k-STM using three kernel configurations: radial basis function (RBF), linear, and a hybrid kernel combining RBF and linear components. Although STM is typically paired with a support vector machine (SVM) as its backbone classifier [11, 12, 19, 21–23], we leveraged the classifier-agnostic nature of STM by adopting four classifiers with fundamentally different structures: logistic regression (LR), SVM, random forest (RF), and multilayer perceptron (MLP). Experimental results demonstrate that k-STM significantly outperforms the conventional STM, particularly when utilizing nonlinear mapping through RBF and hybrid kernels. These findings indicate that k-STM effectively captures nonlinear inter-user variability in sEMG and EEG data, underscoring its potential as a practical and versatile transfer-learning solution for biological signal applications.

## 2 Materials

Four publicly available datasets were used in this study: the MyoDataset [28], NinaProDB5 [29], OpenBMI [30], and BCI Competition IV-2a (BCICIV2a) [31]. The first two datasets consist of sEMG signals, whereas the latter two consist of EEG signals. We primarily followed the pre-processing and feature extraction methodologies described in the corresponding original studies [28–31].

### 2.1 Dataset and Preprocessing

#### 2.1.1 MyoDataset

MyoDataset is a publicly available sEMG dataset collected from 25 healthy participants (aged 20–31, including 7 females). Each participant wore an 8-channel Myo Gesture Control Armband (Thalmic Labs, Canada) on their right forearm, and sEMG signals were recorded at a sampling rate of 200 Hz while performing eight different one-degree-of-freedom (1-DoF) forearm motions. Additionally, participants performed 14 two-degree-of-freedom (2-DoF) combined motions derived from eight basic motions; however, these data were not used in this study. Each motion was recorded for 6 s and repeated five times for each participant.

We conducted the following preprocessing and feature extraction steps. A fifth-order Butterworth high-pass filter with a cutoff frequency of 10 Hz was applied to the raw sEMG signals. Motion onsets were then detected using multi-scale sample entropy, with hyperparameters {*N*_*w*_, *N*_*s*_, *m, r*} adopted from previous studies [32, 33]. From each 6-s trial recording, a 1.5-s segment was extracted, starting from the detected onset. Each segment was divided into 250-ms analysis windows with a 50-ms shift, yielding 26 segments per trial [34].

For the classification, we extracted Huang’s time domain and autoregressive (TDAR) features [35]. Specifically, 11 features were computed per channel: (1) mean absolute value, (2) zero crossing, (3) slope sign changes, (4) waveform length, (5) root mean square, and (6–11) six-order autoregressive coefficients. Consequently, the dimensionality of the feature vector *N*_*d*_ was 88 (8 channels × 11 features).

#### 2.1.2 NinaProDB5

NinaProDB5 is a publicly available sEMG dataset collected from 10 healthy participants (aged 22–34, including 2 females). Each participant wore two Myo Gesture Control Armbands on the right forearm. The distal armband was positioned such that its sensors were in contact with those of the proximal armband, effectively filling the spatial gaps and providing dense sensor coverage. A total of 52 distinct wrist and finger motions were performed by the participants across three experimental tasks (Exercises A, B, and C). In the present study, only the data from Exercise A, comprising 12 finger motion tasks, were analyzed. Each motion was recorded for 5 s and repeated six times per participant.

We conducted the following preprocessing and feature extraction steps. Unlike the MyoDataset, the signals were not filtered. Motion onsets were identified using the “restimulus” variable provided by the dataset publisher [29], and active segments were extracted starting from these onsets. Each segment was divided into 200-ms analysis windows with a 100-ms shift [36].

For classification, we employed Hudgins’ time-domain features [37] and computed four features per channel: (1) mean absolute value, (2) zero crossings, (3) slope sign changes, and (4) waveform length. Consequently, the dimensionality of the feature vector *N*_*d*_ was 64 (2 armbands × 8 channels × 4 features).

#### 2.1.3 OpenBMI

OpenBMI is a publicly available EEG dataset collected from 54 healthy participants (aged 24–35 years, including 25 females). EEG signals were recorded from 62 electrodes placed according to the international extended 10–20 system using a BrainAmp EEG amplifier (Brain Products, Germany) at a sampling rate of 1000 Hz. This study focuses on a motor imagery (MI) experiment involving imagined grasping motions of the left and right hands. Of the four experimental blocks, only the first and second were included in the analysis. Each block consisted of 50 trials for each hand, totaling 100 trials per block. Data from the first block were used to train the classifiers, whereas data from the second block were employed for online testing, with real-time feedback presented to the participants. Each MI trial began with a fixation cross displayed for 3 s, followed by a 4-s cue indicating the imagery task, and concluded with a blank screen lasting 6 s (*±*1.5 s). The dataset providers reported considerable variability in the participants’ abilities to perform MI tasks, identifying only 21 of the original 54 participants as effective performers [30]. Consistent with previous reports on the individual variability in MI task performance [38], we selected 21 participants who successfully completed the tasks for our analysis.

We conducted the following preprocessing and feature extraction steps. EEG signals from 20 electrodes located on the motor cortex (FC5/3/1/2/4/6, C5/3/1/z/2/4/6, CP5/3/1/z/2/4/6) were selected. The original sampling rate of 1000-Hz was downsampled to 100-Hz, and the signals were band-pass filtered between 8 and 30 Hz using a fifth-order Butterworth filter. We extracted 2.5 s segments from 1 to 3.5 s following the onset of the post-cue.

For classification, we extracted the filter bank common spatial patterns (FBCSP) [39]. A filter bank was constructed using six frequency bands (8–12, 12–16, …, 24–28, and 28–30 Hz). For each frequency band, the log-variance features were extracted from the signals projected onto the two highest and two lowest filters of the corresponding spatial filter matrix. Consequently, the dimensionality of the feature vector *N*_*d*_ was 24 (6 bands × 4 filters).

#### 2.1.4 BCICIV2a

BCICIV2a is a publicly available EEG dataset collected from nine healthy participants. Each participant performed four MI tasks: left hand, right hand, foot, and tongue motion imagery. The dataset includes two sessions per participant, each comprising 288 trials, with 72 trials per task. Each trial began with the presentation of a fixation cross and an acoustic warning tone, followed by a visual cue indicating the motor imagery task 2 s later. The visual cue was displayed for 1.25 s, and the participants were instructed to imagine the corresponding motion until the fixation cross disappeared at 6 s post-trial onset.

As the dataset providers did not specify any preprocessing procedures, we adopted the preprocessing steps described by Ang et al. [40]. The signals were bandpass filtered between 4 and 40 Hz using a fifth-order Butterworth filter. We then extracted 2.0-s segments from 0.5 to 2.5 s following the onset of the visual cue.

For classification, we extracted FBCSP using nine frequency bands (4–8, 8–12, …, and 36–40 Hz). Because FBCSP was originally developed for binary classification, we adopted a one-versus-rest (OVR) strategy to address the multiclass classification problem [40]. The log-variance features were extracted from each frequency band of the signals projected onto the two highest and two lowest filters of the corresponding spatial filter matrix. Consequently, the dimensionality of the feature vector *N*_*d*_ was 144 (9 bands × 4 filters × 4 OVR).

### 2.2 Data Splitting

When a participant was designated as a source subject, all of their data were used to pretrain the backbone machine learning models. Conversely, when a participant was designated as the target subject, their data were split into calibration, validation, and test data. The calibration and validation data were used for transfer learning, whereas the test data were reserved for evaluating the transfer learning performance. The data-splitting methodology varied depending on the dataset, as described below.

- **MyoDataset**: The first trial was used for calibration, the second for validation, and the remaining trials for testing.
- **NinaProDB5**: The first, third, and sixth trials were used for calibration; the fourth trial for validation; and the second and fifth trials for testing.
- **OpenBMI**: The first block was randomly partitioned into calibration and validation sets with an 80:20 ratio, and the second block was used entirely for testing.
- **BCICIV2a**: The first session was randomly partitioned into calibration and validation sets with an 80:20 ratio, and the second session was used entirely for testing.

## 3 Methods

### 3.1 Backbone Classifier

We employed four classifiers as backbone machine learning models: logistic regression (LR), support vector machine (SVM), random forest (RF), and multilayer perceptron (MLP). The LR, SVM, and RF models were implemented using scikit-learn, whereas the MLP was implemented using PyTorch. Hyperparameters for each classifier were optimized using Optuna, and 200 trials were conducted per model. The hyperparameter search ranges are summarized in Table 1, where hyperparameters not explicitly listed were set to the default values of the respective libraries. Hyperparameter tuning was performed using 3-fold cross-validation of the data of each source subject.

**Table 1:**
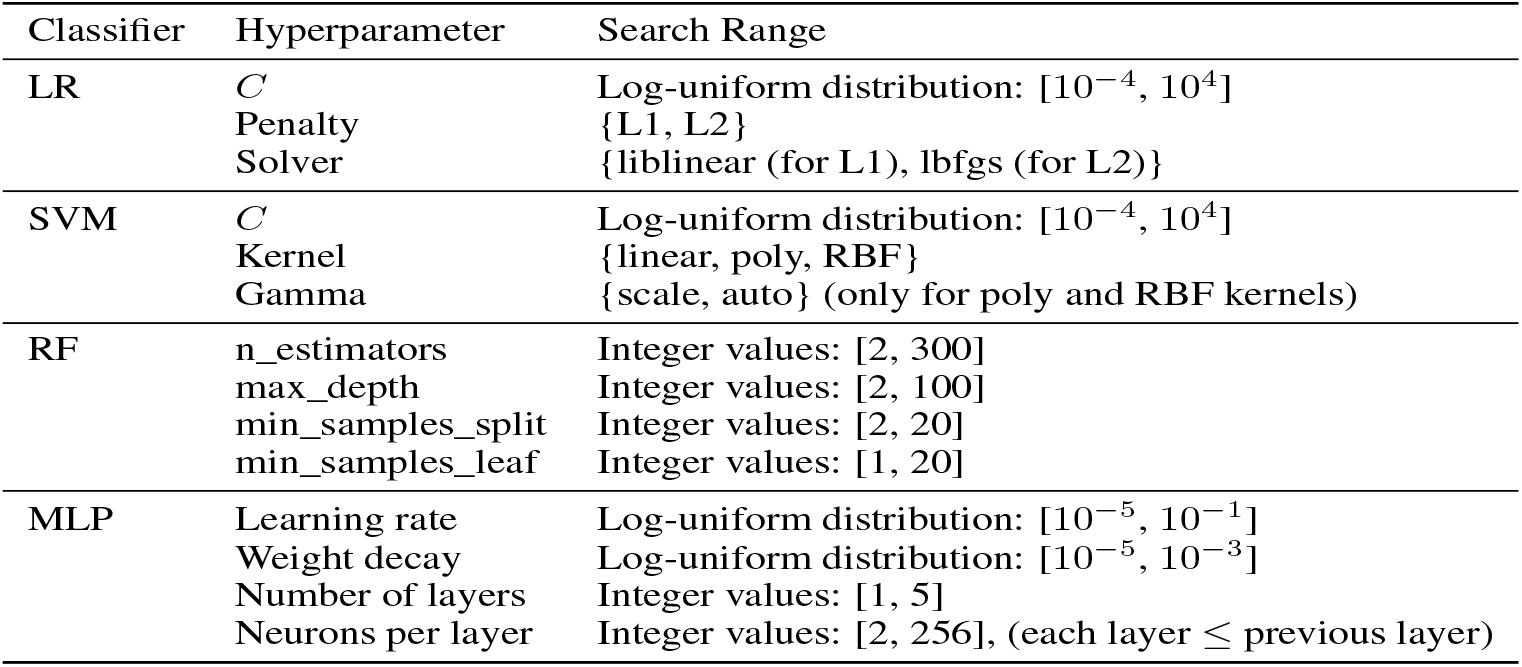
Hyperparameter search ranges used for LR, SVM, RF, and MLP classifiers.

| Classifier | Hyperparameter | Search Range |
| --- | --- | --- |
| LR | $C$ | Log-uniform distribution: $[10^{-4}, 10^4]$ |
| | Penalty | $\{L1, L2\}$ |
| | Solver | $\{\text{liblinear (for L1), lbfgs (for L2)}\}$ |
| SVM | $C$ | Log-uniform distribution: $[10^{-4}, 10^4]$ |
| | Kernel | $\{\text{linear, poly, RBF}\}$ |
| | Gamma | $\{\text{scale, auto}\}$ (only for poly and RBF kernels) |
| RF | n_estimators | Integer values: $[2, 300]$ |
| | max_depth | Integer values: $[2, 100]$ |
| | min_samples_split | Integer values: $[2, 20]$ |
| | min_samples_leaf | Integer values: $[1, 20]$ |
| MLP | Learning rate | Log-uniform distribution: $[10^{-5}, 10^{-1}]$ |
| | Weight decay | Log-uniform distribution: $[10^{-5}, 10^{-3}]$ |
| | Number of layers | Integer values: $[1, 5]$ |
| | Neurons per layer | Integer values: $[2, 256]$, (each layer ≤ previous layer) |

### 3.2 Style Transfer Mapping

The objective of the STM method is to align the data distribution of the target user with that of the source user. To learn the mapping function, representative source data points were first designated as mapping destinations for each calibration sample. Specifically, *k*-means clustering was independently applied to each class within the source data, and the resulting cluster centroids were used as class-specific prototypes. Each calibration sample from the target user was then assigned to the nearest prototype within the same class, thereby determining its corresponding mapping destination. Formally, the *i*-th calibration sample is denoted as 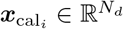, and its corresponding destination prototype is represented as 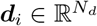. Although previous studies commonly employed 15 clusters [11, 12, 19, 23], we treated the number of clusters as a tunable hyperparameter and explored the values in the set {1, 5, 15, 30}.

Next, we estimated the transformation matrix 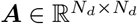 and the bias vector 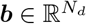 by minimizing the squared differences between each mapped calibration sample and its corresponding prototype. This optimization is formulated as follows:

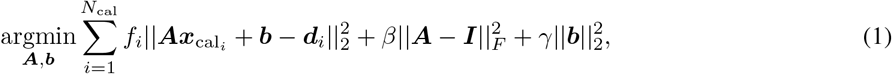

where *f*_*i*_ denotes the confidence value in the range [0, 1], *N*_cal_ is the number of calibration samples, 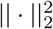 and 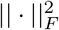 represent the *L*2-norm and Frobenius norm, respectively, and 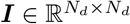 is the identity matrix. The confidence value *f*_*i*_ is a parameter commonly used in semi-supervised learning; in this study, we set *f*_*i*_ = 1 for all the calibration data.

The regularization hyperparameters *β* and *γ* control the degree of transformation, and are defined as follows:

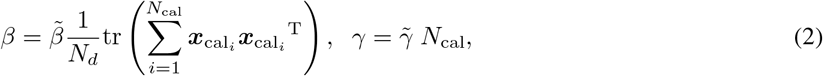

where tr(·) denotes the matrix trace. While previous studies typically explored 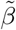 and 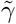 within the range [0, 3] [11,12,19–21], we extended the search range to [0, 10] (see Table 2) because broader ranges may yield improved performance [22]. The hyperparameters 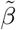 and 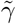 were optimized using Optuna with 200 trials to maximize the classification accuracy of the validation data. Finally, ***A*** and ***b*** were obtained using the following closed-form solutions:

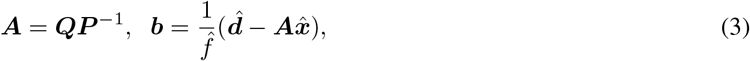

where

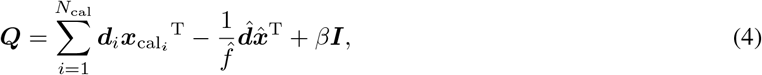

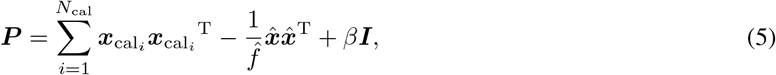

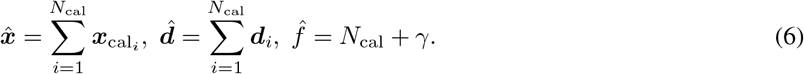

**Table 2:** Hyperparameter search ranges used for STM and k-STM.

| Method | Hyperparameter | Search Range |
| --- | --- | --- |
| STM | $\tilde{\beta}$ | Uniform distribution: $[0, 10]$ |
| | $\tilde{\gamma}$ | Uniform distribution: $[0, 10]$ |
| RBF k-STM | $\sigma$ | Log-uniform distribution: $[10^{-5}, 10^5]$ |
| | $\lambda$ | Log-uniform distribution: $[10^{-5}, 10^5]$ |
| Linear k-STM | $\lambda$ | Log-uniform distribution: $[10^{-5}, 10^5]$ |
| Hybrid k-STM | $\sigma$ | Log-uniform distribution: $[10^{-5}, 10^5]$ |
| | $\lambda$ | Log-uniform distribution: $[10^{-5}, 10^5]$ |
| | $\alpha$ | Uniform distribution: $[0, 1]$ |

Once the optimal mapping parameters ***A*** and ***b*** are obtained, a new unseen data point 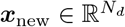 can be projected onto the source distribution using the following affine transformation:

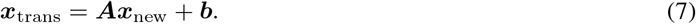

The transformed data ***x***_trans_ are then classified using the source user’s pretrained classifier.

### 3.3 Kernel Style Transfer Mapping

The proposed k-STM extends the linear mapping of the conventional STM to nonlinear mapping and generally follows the same procedure as the original STM method. Prior to executing k-STM, pairs of calibration samples and their corresponding prototypes were prepared as in conventional STM. Let 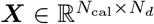 denote the matrix of the calibration samples, and let 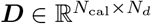 denote the destination matrix containing the corresponding prototypes.

The nonlinear mapping in k-STM is achieved by employing kernel functions, which implicitly project data onto a higher-dimensional feature space. Specifically, we construct a kernel matrix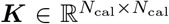, where each entry is computed as 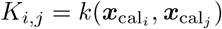, with *k*(·) denoting the chosen kernel function. In this study, we employed three types of kernels: RBF, linear, and hybrid. The RBF kernel is defined as follows.

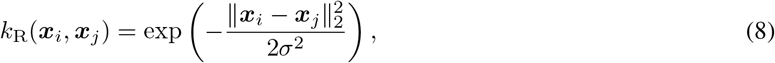

where *σ* denotes kernel bandwidth. The linear kernel is given by the standard inner product

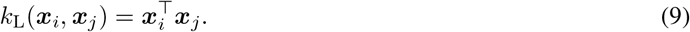

Using this kernel representation, the k-STM learns a weight matrix 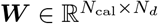 by solving the following optimization problem:

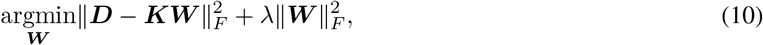

where *λ* is a regularization hyperparameter that controls the trade-off between the fitting accuracy and model complexity. The closed-form solution to this problem is given by

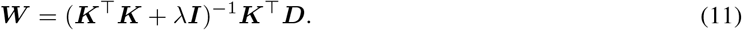

Once the optimal ***W*** is obtained, k-STM can effectively capture the complex nonlinear relationships between the calibration data and their corresponding prototypes.

To map a new unseen data point ***x***_new_ onto the source user’s distribution, we first compute a kernel vector 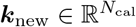, where the *i*th element is given by

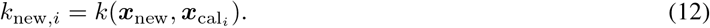

The transformed data were then obtained by applying the learned weight matrix ***W*** :

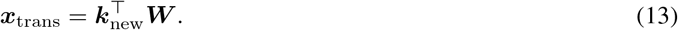

Finally, the transformed data ***x***_trans_ are classified using the source user’s pretrained classifier. Additionally, we introduce a hybrid kernel as a weighted combination of the RBF and linear kernels:

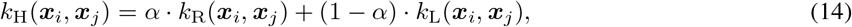

where *α*∈ [0, 1] controls the balance between the RBF and linear components. This hybrid formulation enables the kernel to adapt flexibly between linear and nonlinear mappings, depending on the dataset characteristics. The hyperparameters *σ, λ*, and *α* were optimized using Optuna over 200 trials. The search ranges for the hyperparameters are listed in Table 2. Hereafter, the method is referred to as RBF k-STM when *α* equals 1, linear k-STM when *α* equals 0, and hybrid k-STM when *α* is neither 0 nor 1.

### 3.4 Ensemble Learning

Because STM and k-STM are executed separately for each source subject, an integration step is required to combine their outputs. The mapped data from the target user were classified using each source user’s pretrained classifier. Majority voting based on the average of their predicted probabilities was employed to aggregate the predictions from these multiple classifiers. Although this voting strategy is simple, it has been shown to be effective for transfer-learning tasks involving individual variabilities [41].

## 4 Evaluation

For each dataset, we evaluated the classification accuracy by designating one participant as the target subject and incrementally increasing the number of source subjects from 1 to a maximum of 10. In MyoDataset, for example, when one participant was selected as the target, the remaining 24 participants served as potential source subjects. For a given number of source subjects (e.g., three), we randomly selected distinct source subjects from the available pool and assessed the performances of STM and k-STM. Because evaluating all possible combinations is computationally infeasible, we randomly selected up to 30 non-overlapping combinations for each source-subject count and averaged the resulting accuracy values. This procedure was repeated until each participant was assigned as the target subject.

We performed the following statistical analyses on each dataset to evaluate the effectiveness of the proposed method. First, a three-way repeated-measures analysis of variance (ANOVA) was conducted with the following main factors: (1) transfer learning methods (STM, RBF k-STM, linear k-STM, and hybrid k-STM), (2) number of source users (ranging from 1 to 10), and (3) backbone classifiers (LR, SVM, RF, and MLP). As our primary objective was to assess the effectiveness of the proposed methods, multiple comparisons were performed when a significant main effect was observed for the transfer learning method factor. These comparisons were conducted using paired *t*-tests with Bonferroni correction. The significance level was set at 5

## 5 Results

Figure 3 shows the classification accuracy of each method across all the datasets. Both STM and k-STM exhibited a gradual improvement in accuracy as the number of source subjects increased, regardless of the backbone classifier or dataset. For all datasets, RBF k-STM and hybrid k-STM generally outperformed the conventional STM method. The best-performing configuration for each dataset was as follows: MyoDataset achieved an accuracy of 0.924 using hybrid k-STM with MLP and 10 source subjects; NinaProDB5 achieved 0.711 using hybrid k-STM with LR and nine source subjects; OpenBMI reached 0.865 using RBF k-STM with SVM and seven source subjects; and BCICIV2a obtained 0.670 using linear k-STM with LR and seven source subjects.

**Figure 3:**
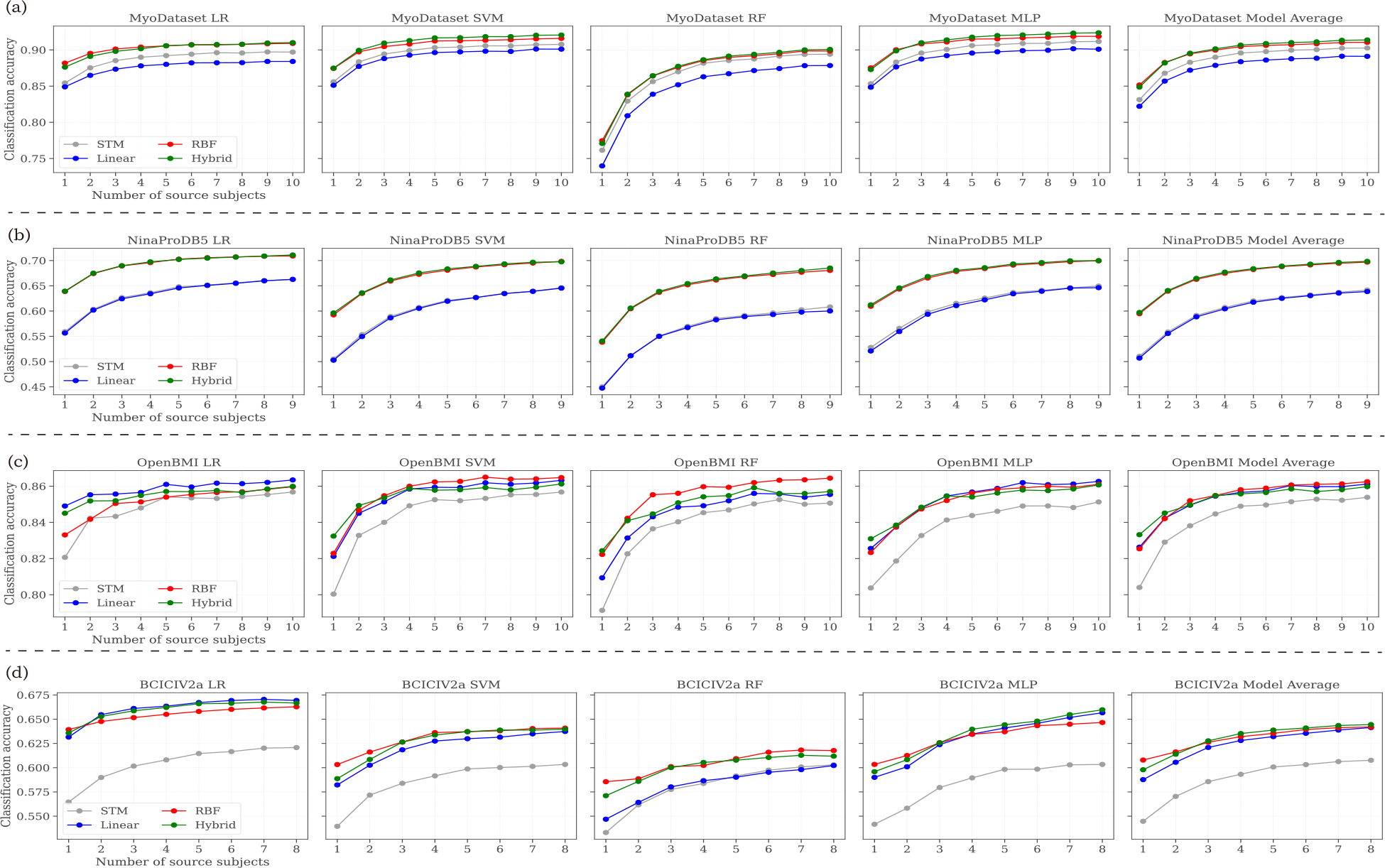
Classification accuracy of STM and k-STM methods across four datasets, shown for varying numbers of source subjects (up to 10, depending on dataset). Each row corresponds to a dataset. Each column corresponds to a backbone classifier, with the fifth column representing the average accuracy across all classifiers.

We conducted a three-way repeated-measures ANOVA for each dataset; the results are presented in Table 3. As shown, all the main effects were statistically significant across all datasets. Furthermore, all two-way interactions were significant, and all three-way interactions were significant for all datasets, except OpenBMI.

**Table 3:** Three-way repeated-measures ANOVA results for each dataset. The factors include *transfer* (transfer learning method: STM, RBF k-STM, Linear k-STM, Hybrid k-STM), *n_src* (number of source subjects), and *model* (backbone classifier: LR, SVM, RF, MLP). Interaction terms (e.g., transfer:n_src) represent the combined effects between factors.

| Dataset | Effect | F-value | Num DF | Den DF | p-value |
| --- | --- | --- | --- | --- | --- |
| MyoDataset | transfer | 23.976 | 3 | 72 | < 0.001 |
| | $n\_src$ | 357.459 | 9 | 216 | < 0.001 |
|  | model | 59.533 | 3 | 72 | < 0.001 |
| | transfer:$n\_src$ | 16.432 | 27 | 648 | < 0.001 |
|  | transfer:model | 3.828 | 9 | 216 | 0.002 |
| | $n\_src$:model | 185.284 | 27 | 648 | < 0.001 |
| | transfer:$n\_src$:model | 2.426 | 81 | 1944 | < 0.001 |
| NinaProDB5 | transfer | 265.041 | 3 | 27 | < 0.001 |
| | $n\_src$ | 492.284 | 8 | 72 | < 0.001 |
|  | model | 64.818 | 3 | 27 | < 0.001 |
| | transfer:$n\_src$ | 26.754 | 24 | 216 | < 0.001 |
|  | transfer:model | 4.737 | 9 | 81 | < 0.001 |
| | $n\_src$:model | 44.192 | 24 | 216 | < 0.001 |
| | transfer:$n\_src$:model | 2.952 | 72 | 648 | < 0.001 |
| OpenBMI | transfer | 4.980 | 3 | 60 | 0.004 |
| | $n\_src$ | 104.122 | 9 | 180 | < 0.001 |
|  | model | 2.868 | 3 | 60 | 0.044 |
| | transfer:$n\_src$ | 6.726 | 27 | 540 | < 0.001 |
|  | transfer:model | 3.711 | 9 | 180 | < 0.001 |
| | $n\_src$:model | 10.495 | 27 | 540 | < 0.001 |
| | transfer:$n\_src$:model | 1.186 | 81 | 1620 | 0.128 |
| BCICIV2a | transfer | 30.758 | 3 | 24 | < 0.001 |
| | $n\_src$ | 61.568 | 7 | 56 | < 0.001 |
|  | model | 19.214 | 3 | 24 | < 0.001 |
| | transfer:$n\_src$ | 11.220 | 21 | 168 | < 0.001 |
|  | transfer:model | 4.888 | 9 | 72 | < 0.001 |
| | $n\_src$:model | 4.337 | 21 | 168 | < 0.001 |
| | transfer:$n\_src$:model | 1.810 | 63 | 504 | < 0.001 |

Because significant main effects of transfer learning were observed for all datasets, we conducted paired *t*-tests with Bonferroni correction. The results are summarized in Table 4. Overall, RBF k-STM and hybrid k-STM significantly outperformed STM across all datasets. For MyoDataset and NinaProDB5, hybrid k-STM achieved the highest performance, significantly surpassing all other methods. For OpenBMI, all k-STM variants significantly outperformed STM, although no significant differences were observed among the kernels. For BCICIV2a, RBF k-STM and hybrid k-STM outperformed the other methods, with no significant differences between them. Taken together, these findings statistically confirm that k-STM, particularly when using nonlinear kernels (e.g., RBF, hybrid), significantly improves classification accuracy compared to conventional STM.

**Table 4:** Bonferroni-corrected paired *t*-test results.

| Dataset | Comparison | t-value | modified p-value |
| --- | --- | --- | --- |
| MyoDataset | STM vs RBF | -14.865 | < 0.001 |
|  | STM vs Linear | 20.841 | < 0.001 |
|  | STM vs Hybrid | -21.644 | < 0.001 |
|  | RBF vs Linear | 32.103 | < 0.001 |
|  | RBF vs Hybrid | -3.427 | 0.004 |
|  | Linear vs Hybrid | -48.877 | < 0.001 |
| NinaProDB5 | STM vs RBF | -55.706 | < 0.001 |
|  | STM vs Linear | 5.103 | < 0.001 |
|  | STM vs Hybrid | -56.693 | < 0.001 |
|  | RBF vs Linear | 57.170 | < 0.001 |
|  | RBF vs Hybrid | -4.262 | < 0.001 |
|  | Linear vs Hybrid | -55.507 | < 0.001 |
| OpenBMI | STM vs RBF | -11.832 | < 0.001 |
|  | STM vs Linear | -15.136 | < 0.001 |
|  | STM vs Hybrid | -14.854 | < 0.001 |
|  | RBF vs Linear | 1.133 | 1.000 |
|  | RBF vs Hybrid | 1.254 | 1.000 |
|  | Linear vs Hybrid | -0.129 | 1.000 |
| BCICIV2a | STM vs RBF | -21.358 | < 0.001 |
|  | STM vs Linear | -19.166 | < 0.001 |
|  | STM vs Hybrid | -26.032 | < 0.001 |
|  | RBF vs Linear | 3.939 | 0.062 |
|  | RBF vs Hybrid | -0.190 | 1.000 |
|  | Linear vs Hybrid | -8.639 | < 0.001 |

### 5.1 Additional Analysis

In this study, we investigated the optimal number of *k*-means clusters (prototypes) for STM and k-STM because previous studies typically fixed this number to 15 without optimization [11, 12, 19, 23]. Because the number of prototypes influences the mapping quality and may affect the classification accuracy, determining the optimal value is crucial for maximizing the performance. To this end, we examined four settings: 1, 5, 15, and 30 clusters, and selected the optimal value based on the validation accuracy for each target-source pair. Figure 4 illustrates the variation in classification accuracy with the number of clusters in the validation dataset. Overall, fewer clusters consistently yielded a higher accuracy for both STM and k-STM, with the best performance observed when using a single prototype. These results suggest that using fewer clusters than previously recommended may improve the classification accuracy.

**Figure 4:**
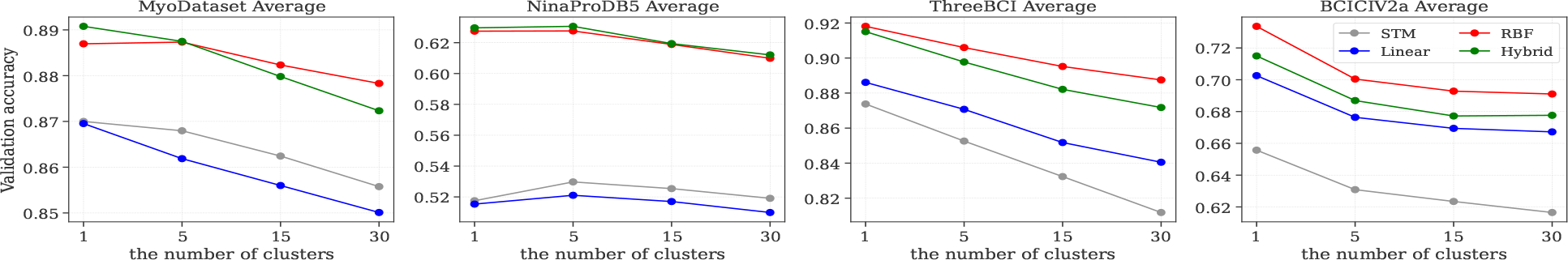
Influence of the number of prototypes (*k*-means clusters) on validation accuracy for each dataset. Each panel shows the averaged results across the four backbone classifiers.

## 6 Discussion

Our statistical analysis demonstrates that transfer learning methods incorporating nonlinear mappings are highly effective for classifying sEMG and EEG signals. In this study, we hypothesized that biological signals exhibit nonlinear intersubject variability and, accordingly, extended the conventional STM framework to a nonlinear variant termed k-STM. The proposed k-STM consistently outperformed the standard STM across all datasets by leveraging nonlinear kernels such as RBF and hybrid kernels. Furthermore, we evaluated a linear kernel in addition to nonlinear kernels (RBF and hybrid). The nonlinear kernels achieved superior classification accuracy across all datasets except the OpenBMI dataset. These findings suggest that the observed performance gains were not merely attributable to the use of kernel methods in general but specifically to the incorporation of nonlinear kernels. Thus, our nonlinear extension of STM effectively mitigates inter-subject variability and enhances classification performance.

The results indicate that hybrid kernels, which balance linear and nonlinear mappings, should be considered the preferred choice for k-STM. Overall, our statistical analyses indicate that the hybrid kernel achieved the highest classification accuracy. However, a more detailed examination revealed that, in certain cases, such as the LR classifier on the OpenBMI and BCICIV2a datasets, the linear kernel performed better. These findings suggest that the optimal kernel selection depends on the specific characteristics of both the datasets and classifiers and that assuming purely linear or purely nonlinear mappings may be overly restrictive. The hybrid kernel addresses this challenge by adaptively balancing the linear and nonlinear components, thereby offering a more flexible and realistic solution. Notably, the effectiveness of combining multiple kernels to enhance classification performance has been demonstrated in previous studies, even beyond the domain of biological signal classification [42], thereby highlighting a promising direction for future research.

In contrast to previous recommendations of using 15 prototypes for each class [11, 12, 19, 23], we found that employing fewer prototypes, particularly a single prototype, led to improved classification accuracy. Although it may seem intuitive to assign each target sample to its nearest prototype among multiple candidates to yield more precise mappings, our results indicate otherwise (see Figure 4). This discrepancy may stem from the risk of overfitting introduced by overly detailed mapping when multiple prototypes are used with limited calibration data. Moreover, using a large number of prototypes can introduce ambiguity in determining the optimal mapping destination, thereby reducing stability and overall accuracy. By contrast, employing fewer prototypes enables target data to be mapped onto more generalized and representative points, ensuring stable correction of individual variabilities and improved classification performance. Therefore, adopting fewer prototypes is recommended for the practical implementation of STM and k-STM.

A key advantage of k-STM is its versatility, which enables seamless integration with classifiers. As articulated by the “No Free Lunch” theorem, no single classifier can consistently achieve optimal performance across all possible problem scenarios [43]. Consequently, transfer learning methods that rely on specific classifiers [15–18] may perform well for certain tasks but exhibit substantially reduced accuracy under different conditions. By contrast, k-STM is a classifier-agnostic transfer learning approach that allows flexible integration with a wide range of classification algorithms, from linear to nonlinear models and from parametric to non-parametric methods. This versatility was empirically validated in our study, where k-STM consistently delivered high performance across classifiers with fundamentally different architectures, including LR, SVM, RF, and MLP.

Furthermore, k-STM can directly leverage insights accumulated from prior STM research. For example, Niu et al. and Kanoga et al. demonstrated that STM is effective as a preprocessing or postprocessing step when combined with other methods [19, 20], and a similar integration is feasible with k-STM. Huang et al. employed STM to fuse multiple source domains into a unified representative source [23]; incorporating k-STM into this process could further enhance the quality of the fused source. Ran et al. proposed the selection of optimal source samples for STM [21], an approach that can also help k-STM mitigate negative transfers by removing detrimental samples. Chen et al. introduced a novel regularization technique to preserve the relationships between the target samples before and after mapping [22], which is particularly beneficial for k-STM. As illustrated in Figure 1, k-STM tends to excessively cluster mapped data around the prototypes owing to its powerful nonlinear mapping capability. Applying Chen et al.’s regularization could provide more effective control over the mapped data distribution, thereby reducing the risk of overfitting. Overall, k-STM inherits and extends these prior advancements, underscoring its potential as a robust and versatile method.

## 7 Conclusions

In this study, we propose k-STM, an advanced nonlinear transfer learning method specifically tailored for cross-subject biological signal classification. Conventional STM, which relies on linear mappings, inherently struggles to capture the complex and nonlinear intersubject variability commonly observed in biological signals, such as EMGs and EEGs. By incorporating nonlinear kernels, including RBF and hybrid kernels, k-STM effectively models nonlinear relationships, leading to significant improvements in classification accuracy. Our evaluation on four public datasets (MyoDataset, NinaProDB5, OpenBMI, and BCICIV2a) using four backbone classifiers (LR, SVM, RF, and MLP) confirmed the superior performance of k-STM over conventional STM, particularly emphasizing the effectiveness of nonlinear kernels. The classifier-agnostic design of k-STM enables seamless integration with a wide range of machine-learning models and existing STM-based methods, highlighting its practicality and adaptability for real-world applications. Future research directions include exploring more sophisticated kernel combinations, extending k-STM to real-time applications, and validating its effectiveness across a broader range of biological signals and classification tasks. Overall, k-STM represents a robust and versatile transfer-learning framework, marking a significant advancement toward practical human–computer interfaces based on biological signals.

## Acknowledgements

This work was partially supported by JST Moonshot R&D (Grant Number JPMJMS2239), by JSPS KAKENHI (Grantin-Aid for Scientific Research (B), Grant Number JP23K28135), and by the Keio University Academic Development Funds for Individual Research.

